# The Cdc42 GEF, Gef1, promotes uniform protein distribution along the actomyosin ring to enable concentric furrowing

**DOI:** 10.1101/381863

**Authors:** Udodirim Onwubiko, Paul J. Mlynarczyk, Bin Wei, Julius Habiyaremye, Amanda Clack, Steven M. Abel, Maitreyi E. Das

**Affiliations:** Department of Biochemistry & Cellular and Molecular Biology, University of Tennessee, Knoxville, TN, USA; Department of Chemical and Biomolecular Engineering, University of Tennessee, Knoxville, TN

**Keywords:** Cdc42, GEF, Cdc15, actomyosin ring, septum, endocytosis

## Abstract

During cytokinesis, fission yeast coordinates actomyosin ring constriction with septum ingression, resulting in concentric furrow formation. Mechanisms coordinating septum ingression with the actomyosin ring remain unclear. We report that cells lacking the Cdc42 activator Gef1, combined with an activated allele of the formin, Cdc12, display non-concentric furrowing. Although cells that furrow non-concentrically display normal actomyosin rings, the scaffold Cdc15 is unevenly distributed along the ring. This suggests that after ring assembly, uniform Cdc15 distribution along the ring drives proper furrow formation. We find that Cdc15 levels at the ring are reduced in the activated *cdc12* mutant, or upon disruption of Arp2/3 complex-dependent endocytic patches. Furthermore, Cdc15 levels in endocytic patches increase in *gef1* mutants. We hypothesize that assembled rings recruit Cdc15 from endocytic patches. Patches with higher Cdc15 levels and slower ring-association rate lead to uneven Cdc15 distribution. Based on this hypothesis we developed a mathematical model that captures experimentally observed Cdc15 distributions along the ring. We propose that, at the ring, Gef1 and endocytic events promote uniform Cdc15 distribution to enable proper septum ingression and concentric furrow formation.

**Summary Statement:** Gef1 and endocytic events at the assembled actomyosin ring facilitate uniform Cdc15 distribution along the ring thus enabling concentric furrow formation.

## INTRODUCTION

Cytokinesis is the final step in cell division. In animal and fungal cells, this involves the constriction of an actomyosin ring which leads to furrow formation (Pollard, 2010). The fission yeast, *Schizosaccharomyces pombe*, has been at the forefront of our understanding of cytokinetic programs based on the actomyosin ring. The cell assembles an actomyosin ring that constricts simultaneously with septum deposition, leading to membrane ingression and furrow formation (Lee et al., 2012; Pollard, 2010). Substantial progress in our understanding of actomyosin ring assembly has been made, largely due to the plenary work by several groups using fission yeast as a model system (Johnson et al., 2012; Lee et al., 2012; Pollard, 2010; Pollard, 2017). Recent reports have also shed light on the mechanistic details of ring constriction in fission yeast (Stachowiak et al., 2014; Thiyagarajan et al., 2015; Zhou et al., 2015). During furrow formation, the constricting actomyosin ring pulls in the membrane adjacent to it to form a physical barrier that separates the two daughter cells. In cell-walled organisms like fission yeasts, the actomyosin ring must overcome high internal turgor pressure to allow for membrane invagination and furrow formation (Proctor et al., 2012). Multiple factors contribute to generate the force necessary furrow formation. First, septum deposition behind the ingressing membrane provides the force necessary to overcome turgor pressure (Proctor et al., 2012). Septum deposition, primarily mediated by the cell-wall synthesizing enzyme Bgs1, promotes furrow formation (Cortes et al., 2002; Cortes et al., 2007; Le Goff et al., 1999; Liu et al., 1999). Second, the type II Myosins, Myo2 and Myp2, are the motors that pull in actin to mediate actomyosin ring constriction (Bezanilla et al., 2000; Mulvihill and Hyams, 2003; Sladewski et al., 2009). These myosins are important for furrow formation even in the presence of an intact septum-building machinery (Laplante et al., 2015). In addition, furrow formation requires membrane expansion at the site of division (Wang et al., 2016). Proper furrow formation requires precise coordination of ring constriction, septum ingression, and membrane expansion that occurs in a circumferentially uniform manner. This causes the furrow to form concentrically towards the central axis of the cell. The mechanistic details that allow proper coordination of ring constriction, septum ingression and membrane expansion are not well understood.

While in most animal cells, the actomyosin ring constricts immediately after assembly, in fission yeast the ring enters a maturation period (Laporte et al., 2010). During the maturation phase, the ring prepares for constriction, septum ingression, and membrane expansion to enable furrow formation (Laporte et al., 2010; Wang et al., 2016). These processes are tightly coordinated such that the furrow forms in a concentric manner, towards the central axis of the cell. Any defect in the coordination of these processes would result in non-concentric furrow formation. It is not clear how these different events are spatio-temporally coordinated during cytokinesis. While the mechanistic events involved in maturation are not clearly understood, several cytokinetic proteins localize to the division site during this phase (Arasada and Pollard, 2014; Bezanilla et al., 2000; Munoz et al., 2013; Ren et al., 2015; Wei et al., 2016). One of the proteins involved in maturation is the F-BAR-domain-containing protein Cdc15, which plays a role in ring formation but is more critical for cytokinetic events after ring assembly (Arasada and Pollard, 2014; Cortes et al., 2015; Martin-Garcia et al., 2014; Ren et al., 2015). Once the ring forms, Cdc15 levels rapidly increase at the division site by an unknown mechanism (Wu and Pollard, 2005). In *cdc15* mutants, while the ring is able to assemble, there are delays in the localization of Bgs1 and defects in ring constriction and septum ingression (Arasada and Pollard, 2014; Cortes et al., 2015). After assembly, the ring also recruits the type II myosin Myp2 in preparation for ring constriction (Bezanilla et al., 2000; Laplante et al., 2015). In addition, coupled membrane trafficking events, such as exocytosis and endocytosis, occur near the division site after ring assembly (Gachet and Hyams, 2005; Wang et al., 2016; Win et al., 2001). Exocytosis mediated by TRAPPII and the exocyst complex promotes delivery of septum-building enzymes, such as Bgs1 and Bgs4, as well as the addition of new membrane for furrow formation (Wang et al., 2016). Soon after ring assembly, proteins involved in vesicle delivery and in endocytosis appear at the division site (Wang et al., 2016). Endocytic proteins such as the Arp2/3 complex, type I myosin Myo1, and fimbrin have been shown to localize to the division site and contribute to cytokinesis (McDonald et al., 2017; Pelham and Chang, 2002; Wang et al., 2016; Wu et al., 2001).

In fission yeast, Cdc42 is activated by two Guanine-nucleotide exchange factors (GEFs), Gef1 and Scd1 (Chang et al., 1994; Coll et al., 2003). Previously, we reported that the small GTPase Cdc42 is activated at the site of cell division in a sequential manner during cytokinesis (Wei et al., 2016). As soon as the actomyosin ring assembles, Gef1 localizes to the ring and activates Cdc42 to promote timely onset of ring constriction and septum ingression. This is followed by Scd1 localization to the ring, where it promotes efficient septum formation. Gef1 is required for timely recruitment of Bgs1 to the ring, while Scd1 recruits Bgs1 to the membrane barrier (Wei et al., 2016). This suggests that Gef1 and Scd1 may function redundantly to localize Bgs1 to the division site. Indeed, while the individual deletion mutants of *gef1* and *scd1* are viable, the *geflscd1* double mutant is inviable (Coll et al., 2003). Gef1 localizes with the constricting ring and is eventually lost once the ring disassembles (Wei et al., 2016). It is not clear how association of Gef1 with the actomyosin ring promotes timely constriction and septum ingression.

Here we show that Gef1 is required for concentric furrow formation. We find that a *gef1* deletion mutation combined with an activated allele of the formin *cdc12* results in non-concentric membrane furrowing. In these mutants, the rate of actomyosin ring constriction is not consistent along the circumference of the ring, with certain regions constricting faster than others. While actomyosin ring assembly is normal in these mutants, Cdc15 is distributed unevenly along the ring. Regions of the ring with a larger fraction of Cdc15 furrow faster than regions containing lower levels of Cdc15. Cdc15 dynamics, measured by fluorescence recovery after photo bleaching, at the ring are slower in the activated formin mutants. In addition, these mutants display lower levels of Cdc15 at the ring after ring assembly. We show that after assembly, disrupting Arp2/3-complex-mediated endocytic actin patches results in lower levels of Cdc15 at the ring. Our data show that Gef1 limits the level of Cdc15 in endocytic patches in a Cdc42-dependent manner. We hypothesize that after ring assembly, Cdc15 is recruited to the division site via endocytic actin patches. Slower dynamics of patch recruitment and increased amount of Cdc15 in individual patches lead to its uneven distribution along the ring. Based on this hypothesis, we develop a mathematical model that recapitulates our experimental observations regarding Cdc15 distribution along the actomyosin ring. These findings suggest that Gef1-mediated Cdc42 activation regulates Cdc15 organization at the ring after assembly to promote concentric furrow formation.

## RESULTS

### Cdc42 GEF Gef1 promotes robust actomyosin ring constriction

We have previously shown that Gef1 is required for timely onset of ring constriction and septum ingression, and that cells lacking *gef1* display fewer non-medial actin cables (Wei et al., 2016). We asked whether the decrease in non-medial actin cables results in the delayed onset of ring constriction we observe in *gef1* mutants. The formin Cdc12 contributes to non-medial actin cable formation during cytokinesis. We hypothesized that the activated allele of the formin *cdc12* would restore non-medial actin cables in *gef1Δ* mutants and rescue the constriction defect. To test this, we first observed the actin cytoskeleton in cells stained with Alexa Fluor Phalloidin. We found that the activated formin mutant *cdc12Δ503* increased the number of actin cables in *gef1Δ* mutants (Fig.1A). Non-medial actin cables were counted in the ring phase of cytokinesis. *gef1+cdc12+* cells displayed a mean value of 3 non-medial actin cables per cell, while *gef1Δ* mutants showed a mean value of 2 cables per cell. Both *cdc12Δ503* and *gef1Δcdc12Δ503* mutants showed an increase in non-medial actin cables with a mean value of 4 and 5 actin cables per cell, respectively (p=0.01 and <0.0001respectively, Fig.1A). In addition to an increase in the number of actin cables, *cdc12Δ503* and *gef1Δcdc12Δ503* mutants also displayed disorganized endocytic actin patches (Fig. 1B). The actin patches in these mutants are not restricted to the cell ends or the site of cell division, as is observed in *gef1+cdc12+* and *gef1Δ* cells (Fig. 1B).

**Figure 1.**
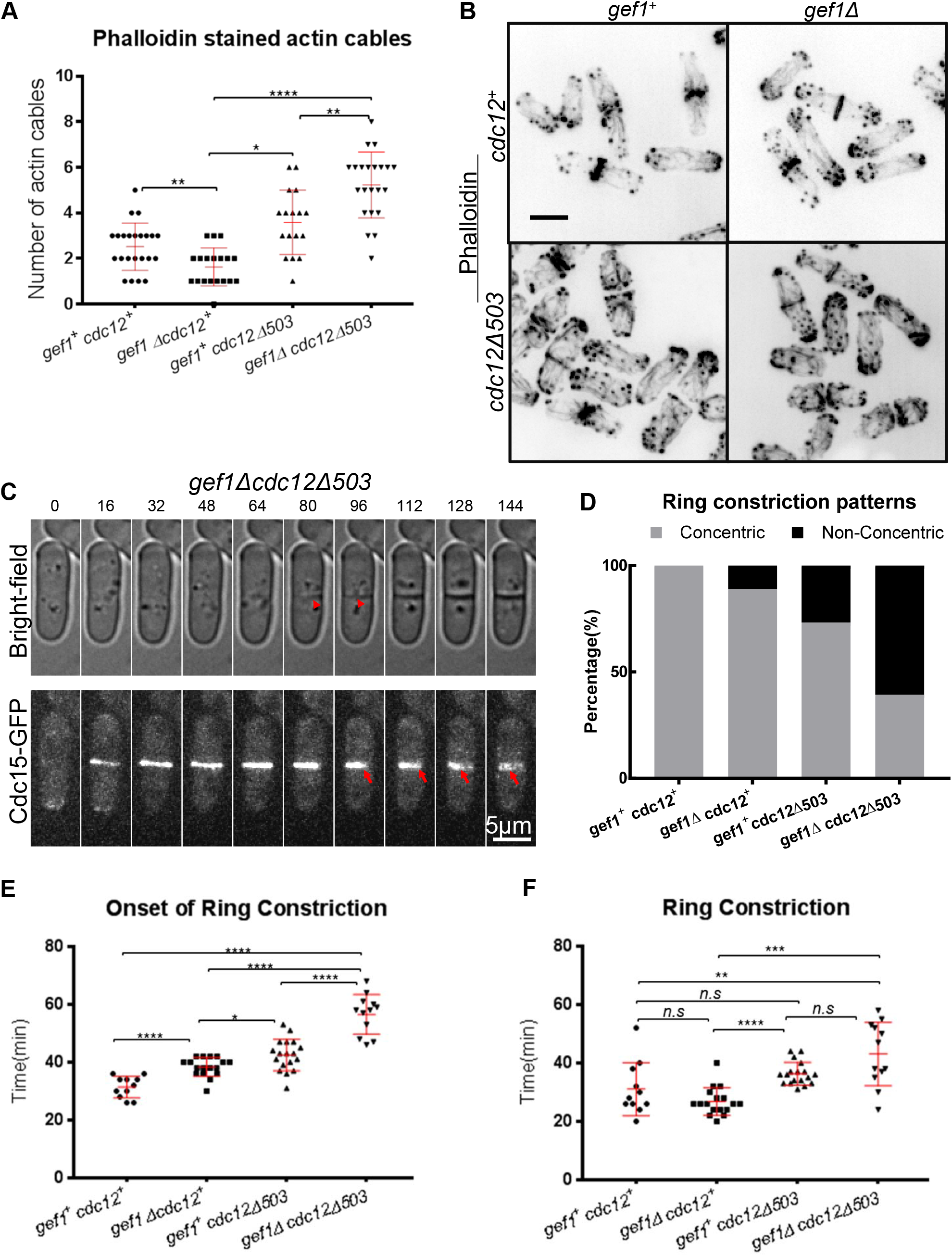
Gefl promotes robust actomyosin ring constriction. **(A)**. Quantification of Phalloidin stained non-medial actin cables in strains as indicated. **(B)**. Phalloidin staining of actin cytoskeleton organization in strains as indicated. **(C)**. Time-lapse image showing non-concentric furrow formation in a *gef1Δ cdc12Δ503* mutant at16 minute intervals. Red arrowheads depict the septum in the bright-field images. Red arrows depict Cdc15-GFP labelled actomyosin ring. **(D)**. Quantification of non-concentric furrow formation in *gef1*^+^*cdc12*^+^ (n=17), *gef1Δ* (n=27), *cdc12Δ503* (n=41) and *gef1 Δcdc12Δ503* (n=28) cells. **(E)**. Quantification of onset of constriction of Cdc15-GFP labelled rings in the indicated strains (n≥11 cells). **(F).** Quantification of duration of constriction of Cdc15-GFP labelled rings in the indicated strains (n≥11 cells). *n.s-not significant,* p≤0.05, ** p<0.01, ***p<0.001, ****p<0.0001*; *error bars, standard deviation; Scale Bar, 5μm*.

Next we analyzed cytokinetic events in *gef1^+^cdc12^+^, gef1Δ, cdc12Δ503* and *gef1Δcdc12Δ503* cells. We used the F-BAR-domain-containing protein Cdc15-GFP as an actomyosin ring marker and Sad1-mCherry as a spindle pole body marker to analyze different cytokinetic events. Contrary to our hypothesis, we found that the severity of ring constriction defects was increased in *gef1Δ* mutants expressing the activated formin mutant *cdc12Δ503. gef1Δcdc12Δ503* cells displayed a unique phenotype in which the cleavage furrow failed to form in a concentric manner. In these mutants, the ring marked by the F-BAR protein Cdc15-GFP tends to constrict to one side of the cell instead of the central axis. As a result, the furrow forms non-concentrically from one side of the cell to the opposite side as observed by bright field microscopy (Fig.1C). In *gef1+cdc12+cells*, ring constriction and septum ingression are coordinated and as a result the furrow forms concentrically towards the central axis of the cell. Non-concentric furrowing was observed in a small fraction of *gef1Δ* (11%) and *cdc12Δ503* mutants (27%; Fig. 1D). However, in *gef1Δcdc12Δ503* mutants, non-concentric furrowing occurred in 61% of cells. These data suggest that Gef1 is required for robust cytokinesis and contributes to the process that ensures concentric furrow formation.

To understand the nature of the defect leading to non-concentric furrow formation, we analyzed cytokinetic events in *gef1Δcdc12Δ503* mutants in more detail. In both *gef1+cdc12+and gef1Δ* cells, ring formation occurs about 11 minutes after the spindle pole body separates, and was comparable to *cdc12Δ503* mutants that assembled the actomyosin ring at 12 minutes after spindle pole body separation (Supplemental material Fig. S1A). In *gef1Δcdc12Δ503* mutants, we observed prolonged actomyosin ring assembly, which lasted about 15 minutes (Supplemental material Fig. S1A). In addition, *gef1Δcdc12Δ503* mutants displayed a delay in the onset of ring constriction. In *gef1^+^cdc12*^+^cells, onset of ring constriction occurs about 31 minutes after spindle pole body separation (p<0.0001). As reported earlier for *gef1Δ* mutants, this event is delayed by 38 minutes (Fig1E). Onset of ring constriction is further delayed in *cdc12Δ503* and *gef1Δcdc12Δ503* mutants to about 43 and 57 minutes respectively (p<0.0001). Both *gef1Δ* and *gef1+cdc12+* cells completed ring constriction in ~27 minutes (Fig. 1F). In contrast, in both *cdc12Δ503* and *gef1Δcdc12Δ503* mutants, the duration of ring constriction was decreased and lasted on average for 36 and 46 minutes, respectively (p=0.09, and p=0.008) (Fig. 1F). These data demonstrate that in *gef1Δcdc12Δ503* mutants, ring constriction is impaired and this likely results in non-concentric furrow formation.

### Non-concentric ring constriction was observed in cells with uneven distribution of the F-BAR protein Cdc15

Non-concentric furrow formation is a likely outcome of loss of coordination of ring constriction and septum ingression. To define the factors that ensure concentric furrowing, we analyzed the constriction defects in *gef1Δcdc12Δ503* mutants. We followed constriction patterns of Cdc15-GFP-labeled actomyosin rings in *gef1Δcdc12Δ503* mutants over time to generate kymographs (Fig. 2A,a-d). We found that in *gef1Δcdc12Δ503* mutants, the ring tends to constrict away from the central axis of the cell, as represented by the red dashed line, indicating non-concentric constriction (Fig. 2A,a-d). In these mutants, some sections of the ring constricted faster compared to the rest, suggesting that non-concentric furrowing occurred due to an irregular constriction rate along the ring. Normally, before the onset of ring constriction, protein distribution along the ring appears uniform. We measured the intensity of Cdc15-GFP at the two outer edges of the z-projected rings in the kymographs just before the onset of constriction. We calculated the fraction of Cdc15-GFP intensity at each edge in comparison to the total at each ring. In the kymographs, sections of the ring that displayed faster rates of constriction also displayed a higher fraction (0.64) of Cdc15-GFP intensity prior to the onset of constriction (Fig. 2B). In comparison, sections of the ring displaying a slower rate of constriction had a smaller fraction (0.36) of Cdc15-GFP intensity (Fig. 2B). The fraction of *gef1Δcdc12Δ503* rings that displayed concentric furrow formation showed an equivalent distribution of Cdc15 intensity along the two edges of the z-projected ring. (Fig. 2A,d, B). Thus, furrows that constricted in a non-concentric manner also exhibited irregular distribution of Cdc15-GFP along the ring, and sections of the ring with increased protein levels displayed faster constriction.

**Figure 2.**
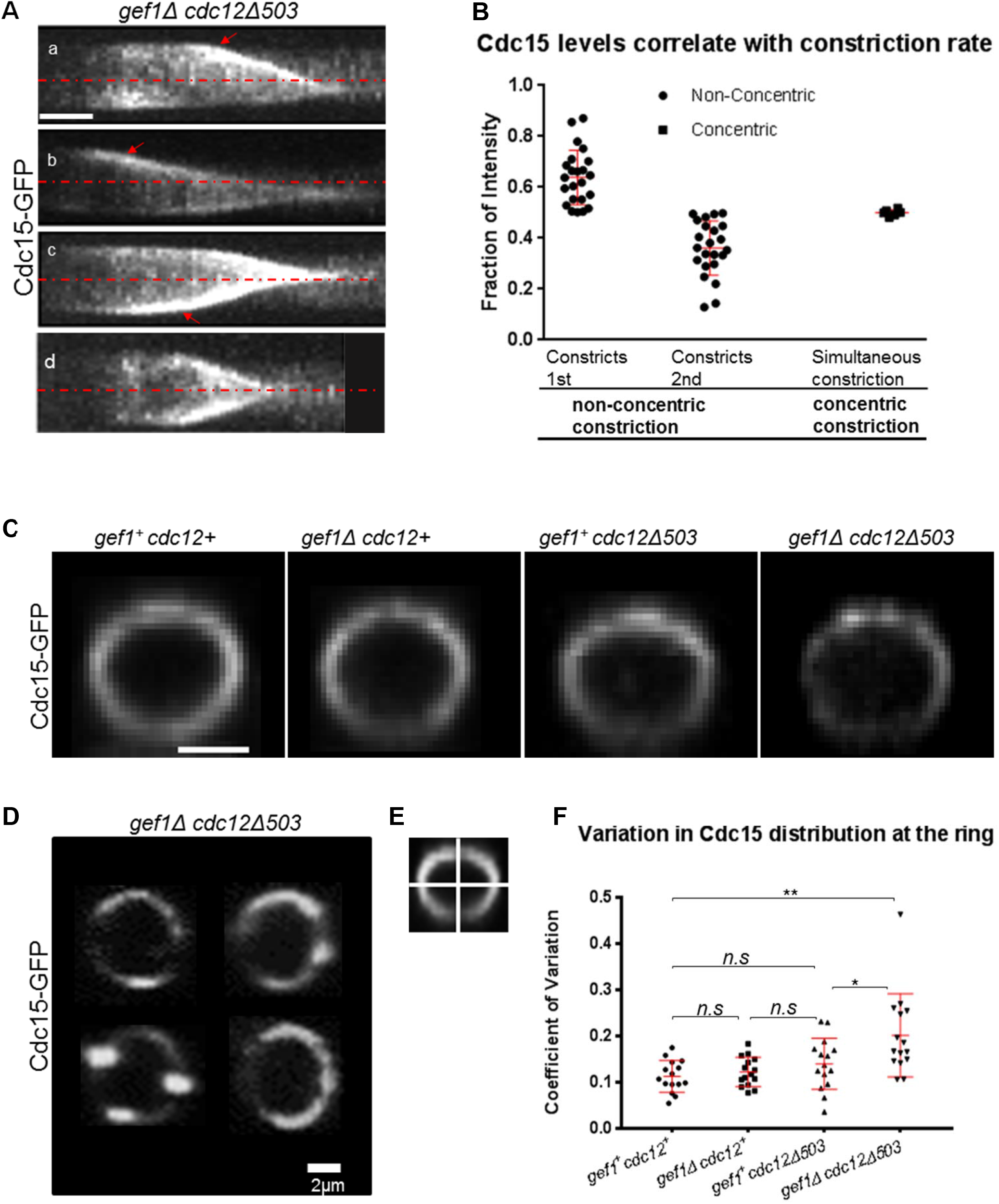
Cells displaying non-concentric furrow formation exhibit uneven Cdc15 distribution along the ring. **(A)**. Kymographs from time-lapse movies showing constriction of Cdc15-GFP labelled rings in *gef1Δcdc12Δ503* cells. Red dashed lines indicate central axis of the cells. Red arrows show regions of the ring constricting at a faster rate. Scale Bar, 4μm. **(B).** Quantification of the fraction of Cdc15-GFP intensity at different regions of the ring immediately prior to the onset of constriction with the corresponding constriction pattern (n=25cells). **(C).** 3D reconstructed Cdc15-GFP labelled actomyosin ring of the indicated strains. **(D).** Examples of Cdc15-GFP distribution along the ring in *gef1Δcdc12Δ503* cells. Scale bar, 2μm. **(E).** Illustration of actomyosin ring quadrants used for analyzing the Coefficient of Variation of Cdc15-GFP distribution. **(F).** Quantification of the Coefficient of Variation of Cdc15-GFP distribution in the indicated strains(n=15 cells). *n.s.-not significant*, \**p=0.033*, \*\**p=0.002*; *Error bars, standard deviation*.

This suggested that Cdc15 is irregularly distributed along the ring in *gef1Δcdc12Δ503* mutants, when compared to *gef1^+^cdc12^+^, gef1Δ*, and *cdc12Δ503* cells. To test this, we divided the rings into four equal quadrants and measured the intensity of Cdc15-GFP in each quadrant, as previously described (Wei et al., 2017). Next, we computed the coefficient of variation (CV) of the intensities of the quadrants of each ring. A higher coefficient of variation indicates more irregular distribution of proteins along the ring. We find that prior to the onset of constriction; the coefficient of variation in assembled rings was comparable in *gef1^+^cdc12^+^, gef1Δ*, and *cdc12Δ503* cells (Fig. 2C,F). However, it was higher in *gef1Δcdc12Δ503* mutants (p=0.002; Fig. 2C, D, F) suggesting increased variation in Cdc15-GFP distribution in these rings. Thus, cells that furrow non-concentrically display irregular distribution of Cdc15-GFP along the ring, and sections of the ring with higher Cdc15-GFP tend to constrict faster.

It is possible that irregular distribution of Cdc15-GFP along the ring in *gef1Δcdc12Δ503* mutants was due to defects in ring assembly or a structural anomaly in the actomyosin ring. To test this, we analyzed actomyosin rings in *gef1^+^cdc12^+^, gef1Δ, cdc12Δ503*, and *gef1Δcdc12Δ503* cells expressing LifeAct-3xmCherry as an actin marker. Under our imaging conditions we did not find any structural anomaly in the actomyosin ring in these cells (Fig. 3A). Further we analyzed the distribution of actin along the ring in these cells as done previously for Cdc15-GFP. We measured the intensity of LifeAct-3xmCherry in the four quadrants of the actomyosin ring and computed the coefficient of variation. We found that coefficient of variation is comparable in *gef1^+^cdc12^+^ gef1Δ, cdc12Δ503*, and *gef1Δcdc12Δ503* cells (Fig. 3C). These findings suggest *gef1Δcdc12Δ503* mutants show no defects in the organization of actin along the actomyosin ring as compared to the control cells. The actomyosin ring contains the type II myosin light chain Rlc1 that is required for ring assembly. Similar to our findings with actin, the coefficient of variation of Rlc1-tdTomato distribution along the rings were comparable in *gef1^+^cdc12^+^, gef1Δ, cdc12Δ503* and *gef1Δcdc12Δ503* cells (Fig. 3B, D). This presents additional evidence that the assembled actomyosin ring in *gef1Δcdc12Δ503* cells have no discernable structural defects.

**Figure 3.**
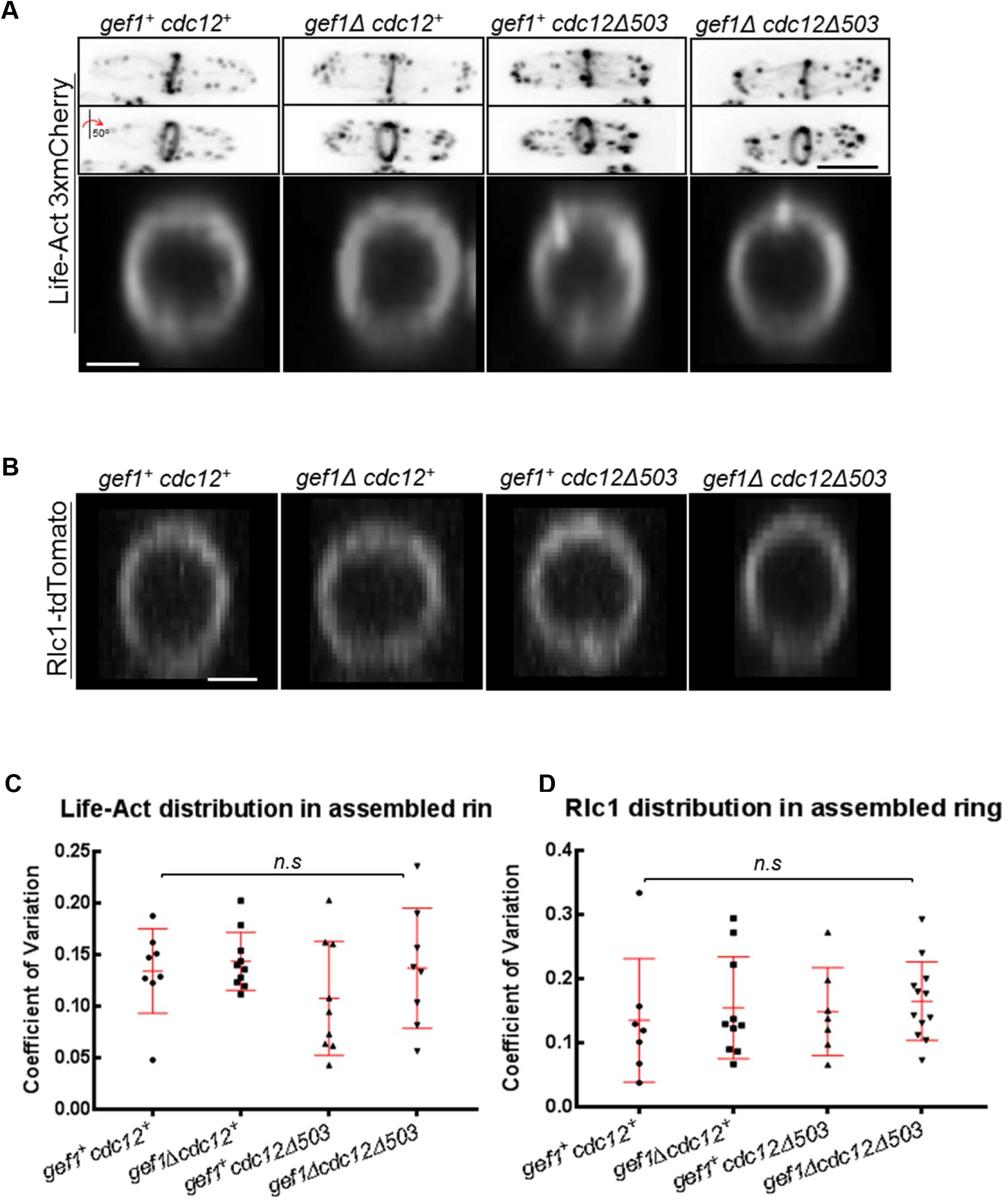
Actin and myosin in the rings appear normal in mutants displaying non-concentric furrow formation. **(A).** 3D projections of cells expressing Life-Act 3xmCherry in indicated rings. Middle panel shows cells rotated by an angle of 50°. Lower panel shows 3D-reconstructed rings labelled with Life-Act-3xmCherry in indicated strains. **(B).** 3D-reconstructed rings labelled with type II myosin light chain Rlc1-tdTomato in the strains as indicated. **(C).** Quantification of the Coefficient of Variation of Life-Act-3xmCherry distribution along the ring in the indicated strains. (n=9) **(D).** Quantification of the Coefficient of Variation of Rlc1-tdTomato distribution along the ring in the indicated strains. (n≥7). *n.s.-not significant, error bars, standard deviation; Scale bar for cells, 5μm; Scale bar for rings, 2μm*.

Cdc15 appears at the precursor nodes and helps recruit Cdc12 thus contributing to ring formation (Carnahan and Gould, 2003; Willet et al., 2015). The *cdc12Δ503* allele is a truncated mutant that does not have the first 503 amino acids of Cdc12 (Coffman et al., 2013). This truncated mutant does not contain the Cdc15 interacting site. Irregular distribution of Cdc15 could be explained by the lack of this interaction. To test this, we analyzed Cdc15-GFP distribution along the ring in *cdc12P31A* mutants in a *gef1Δ* background. The *cdc12P31A* mutant allele lacks only the Cdc15 interacting site and fails to bind this protein. We found that the coefficient of variation of Cdc15-GFP in *gef1Δcdc12P31A* rings was comparable to that of the rings in *gef1^+^cdc12^+^, gef1Δ*, and *cdc12P31A* cells (Supplementary material Fig. S1). This indicated that the irregularity in Cdc15 distribution at the ring in *gef1Δcdc12Δ503* mutants is not due to the lack of Cdc12-Cdc15 protein interaction.

To understand how Gef1 regulates Cdc15 organization in the ring we looked at known downstream targets of Gef1. Gef1 activates Cdc42 which in turn activates the PAK kinase Pak1. Cdc15 is a highly phosphorylated protein and when de-phosphorylated it forms oligomers via the BAR domain and establishes proper interaction with the membrane. We asked if irregularities in Cdc15 distribution along the ring are due to absence of Pak1 dependent phosphorylation in *gef1Δcdc12Δ503* mutants. To test this, we analyzed Cdc15 distribution along the ring in the *pak1* mutant allele *orb2-34* and *orb2-34cdc12Δ503* mutant. The coefficient of variation for Cdc15 distribution along the ring was not altered in *orb2-34* or *orb2-34cdc12Δ503* mutants in comparison to *orb2+cdc12+* cells (Supplemental material Fig.S1C). This indicates that irregularities in Cdc15 distribution along the ring in *gef1Δcdc12Δ503* mutants are not due to lack of Pak1 kinase activity. This raises the question: Why is Cdc15-GFP distribution along the ring disrupted in *gef1Δcdc12Δ503* cells? Could events that occur after ring assembly determine uniform distribution of Cdc15 along the ring? Irregular distribution of Cdc15 along the ring likely results in non-concentric furrow formation. Understanding what enables uniform Cdc15 distribution along the ring will shed light on what causes non-concentric furrow formation and how ring constriction and septum ingression are coordinated. To investigate this, we looked into how Cdc15 is recruited and loaded onto the ring.

### Activated *cdc12* mutants display reduced Cdc15 levels in the ring

Cdc15 localizes to the precursor nodes prior to ring assembly to promote ring formation. After assembly, Cdc15 levels rapidly increase at the ring until the onset of constriction. Previous reports have shown that Cdc15 is a highly abundant protein with fast dynamics at the ring (Laporte et al., 2011; Roberts-Galbraith et al., 2010). To understand how Cdc15-GFP is distributed along the ring we first investigated its dynamics at the actomyosin ring. We performed fluorescence recovery after photo bleaching (FRAP) of Cdc15-GFP at rings during maturation in *gef1^+^cdc12^+^, gef1Δ, cdc12Δ503*, and *gef1Δcdc12Δ503* cells (Fig. 4A). Cdc15-GFP recovery for *gef1Δ* mutants was comparable to that of *gef1^+^cdc12^+^cells* with a t_1/2_ of 17 and 16 seconds, respectively (Fig. 4B). In contrast, recovery was slower in *cdc12Δ503* and *gef1Δcdc12Δ503* cells with a t_1/2_ of 24 and 30 seconds respectively (Fig. 4A, B). We did not see any change in the total recovery of Cdc15-GFP in the cells (Fig. 4A). Hence after ring assembly in *cdc12Δ503* and *gef1Δcdc12Δ503* cells, additional recruitment of Cdc15-GFP to the ring is inefficient compared to *gef1^+^cdc12^+^ and gef1Δcdc12*^+^ cells.

**Figure 4.**
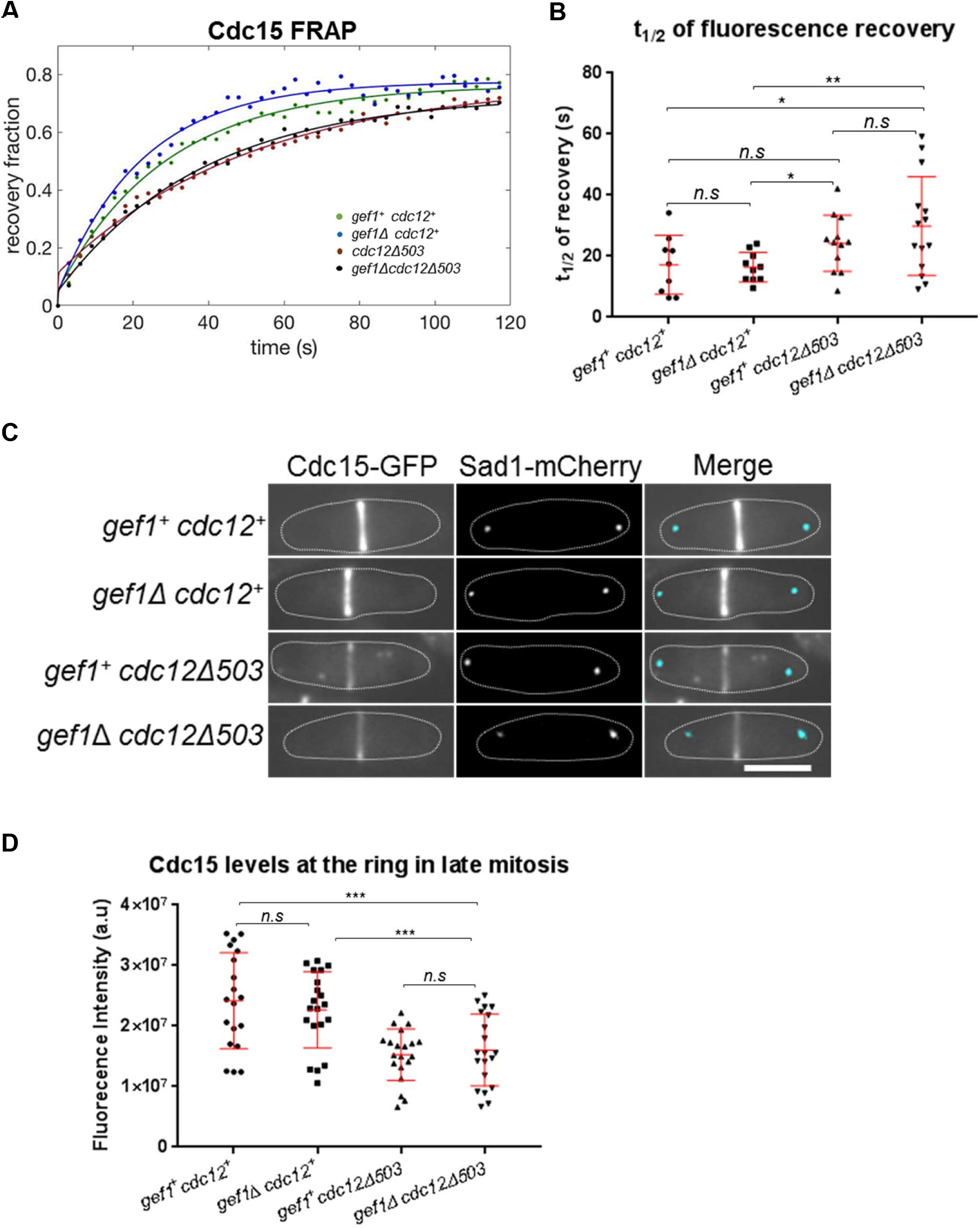
Cdc15 dynamics are slower in activated formin mutants. **(A)**. Quantification of fluorescence recovery after photo-bleaching of Cdc15-GFP at the ring during maturation in the strains as indicated. **(B)** Quantification of the t1/2 of fluorescence recovery after photo-bleaching of Cdc15-GFP in the strains as indicated (n≥9cells). **(C).** Sum projections of images of cells in anaphase B, expressing Cdc15-GFP. The spindle pole bodies are labelled with Sad1-mCherry. **(D).** Quantification of fluorescence intensity of Cdc15-GFP at the actomyosin ring in cells as indicated above, (n=20 cells). *n.s-not statistically significant*; \**p≤0.05; **p<0.01; ***p<0.001; error bar, standard deviation; Scale bar, 5μm*.

Slower protein dynamics as observed by FRAP is typically associated with weaker protein recruitment. We hypothesize that *cdc12Δ503* and *gef1Δcdc12Δ503* mutants show weaker Cdc15 recruitment to the ring. To assess this, we compared the intensity of Cdc15-GFP along the ring during maturation in *cdc12Δ503* and *gef1Δcdc12Δ503* cells to that of *gef1*^+^*cdc12*^+^ and *gef1Δ* cells. Cells in Anaphase B have rings in the maturation phase, and are about to begin constriction (Wei et al., 2017; Wu et al., 2003). Cdc15-GFP intensity was measured in the rings of cells undergoing Anaphase B as determined by the distance of the spindle pole body marker Sad1-mCherry (Fig. 4C). We find that Cdc15-GFP fluorescence intensities at the ring during Anaphase B in *gef1Δ* mutants are similar to those in *gef1*^+^*cdc12*^+^*cells* (Fig. 4C,D). In contrast, the Cdc15-GFP fluorescence intensity at the ring was decreased in both *cdc12Δ503* by ~37% and in *gef1Δcdc12Δ503* by ~34% cells during Anaphase B (p≤0.001; Fig. 4D). These findings are in agreement with the slower Cdc15 dynamics in the activated formin mutant reported above. Thus, our data suggest that Cdc15 localization to the actomyosin ring during maturation is impaired in the activated formin mutant.

### Cdc15 recruited to the assembled actomyosin ring requires membrane trafficking events

How Cdc15 is recruited to the assembled actomyosin ring after assembly remains unclear. We find that in the activated formin mutant, Cdc15 localization to the ring is impaired. As shown above the activated formin mutants display disorganized actin cytoskeleton with increased actin cables (Fig.1A,B). Thus we asked if Cdc15 localization to the ring during maturation is actin dependent. Previous reports have shown that Cdc15 localizes to endocytic patches at the cortex and is involved in endocytosis (Arasada and Pollard, 2011; Carnahan and Gould, 2003). Endocytosis at the cell division site initiates during maturation after actomyosin ring assembly (Wang et al., 2016). Could Cdc15 be recruited to the division site via endocytic patches? Endocytic patches require the actin nucleator Arp2/3 complex that forms branched actin filaments. To test whether Cdc15 recruitment is affected by endocytosis, we treated Cdc15-GFP expressing cells with the Arp2/3 complex inhibitor CK666. CK666 treatment disrupted endocytic actin patches as determined by the loss of actin patches in LifeAct-3xmCherry labelled cells (Supplementary material Fig.2A). Similarly, Cdc15-GFP labelled patches at the cortex were also lost in cells within 5 mins of CK666 treatment (Fig. 5A). Treatment with CK666 did not block actomyosin ring assembly as determined by Rlc1-tdTomato signal at the ring (Fig. 5E). Furthermore, CK666 treatment did not disrupt Cdc15 recruitment to the ring during assembly (Fig. 5B). In DMSO treated control cells, Cdc15 intensity levels after ring assembly continued to rise with a 3-fold increase over a 10-minute period similar to previous reports (Fig. 5C, D) (Wu and Pollard, 2005). However, after assembly, the ring in CK666 treated cells did not show any increase in the intensity of Cdc15-GFP (Fig. 5C,D). Our data suggests that Cdc15 recruitment to the ring depends on two independent mechanisms. During ring assembly, Cdc15 is recruited via precursor nodes (Laporte et al., 2011). After ring assembly, localization of Cdc15 to the actomyosin ring depends on Arp2/3 dependent actin nucleation.

**Figure 5.**
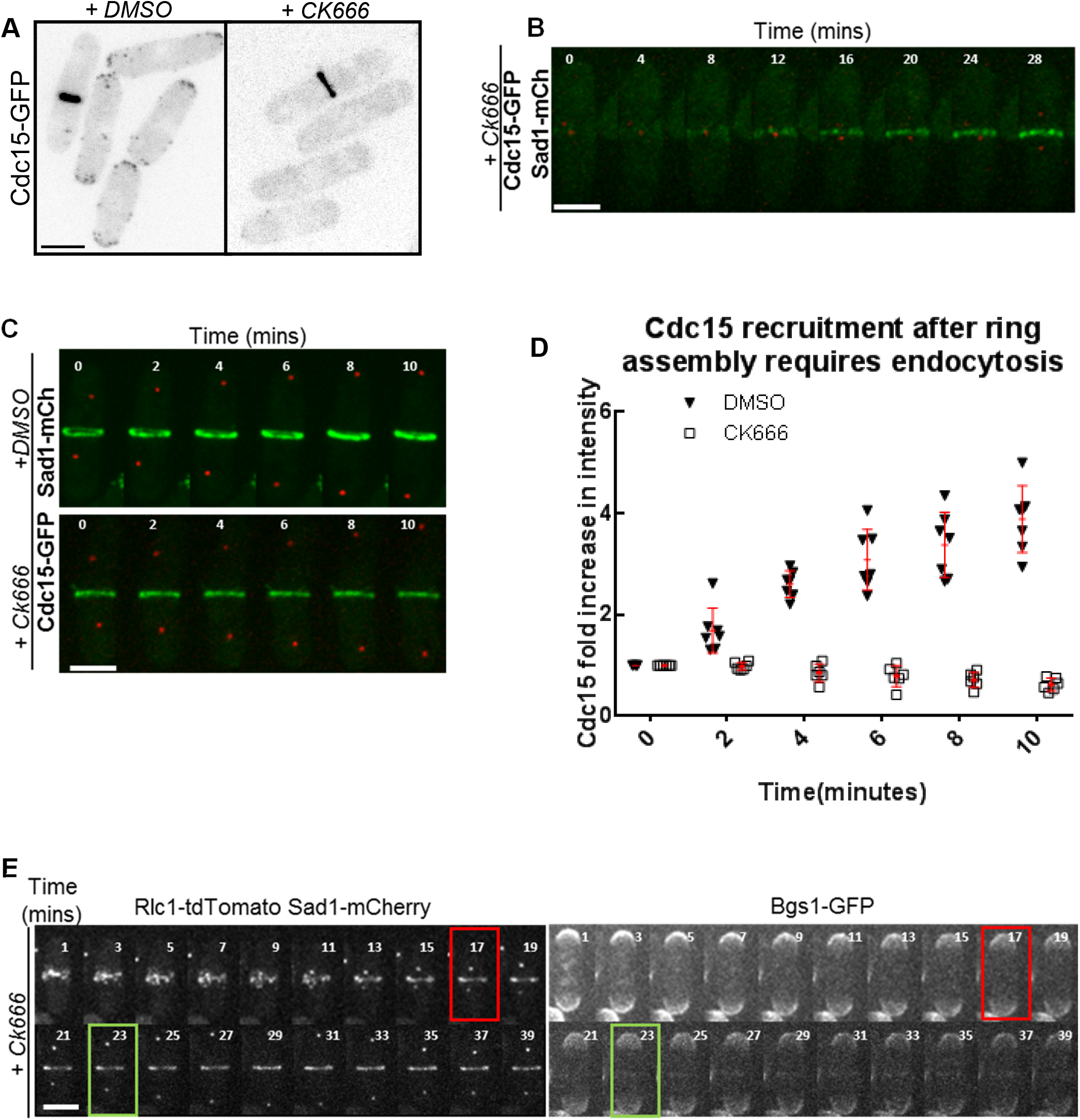
Cdc15 recruited to the assembled actomyosin ring requires Arp2/3 complex mediated events. **(A)**. Cdc15-GFP labelled cortical patches in cells treated with DMSO or 100μM CK666 for 5 minutes. **(B).** Time-lapse images showing Cdc15-GFP recruitment during ring assembly in cells treated with 100μM CK666. Spindle pole bodies are labeled with Sad1-mCherry. **(C).** Time-lapse images of assembled rings in cells expressing Cdc15-GFP treated with DMSO or 100μM CK666. Spindle pole bodies are labeled with Sad1-mCherry. **(D).** Quantification of fold increase in Cdc15-GFP intensities over a 10 minute time interval in cells treated with DMSO treated or 100μM CK666, (n≥7cells). **(E).** Time-lapse images of 100μM CK666 treated cells expressing Bgs1-GFP, Rlc1-tdTomato and Sad1-mCherry during ring assembly and maturation. Red box indicates an assembled actomyosin ring, green box indicates when Bgs1-GFP first appears at the ring. *Error bars, standard deviation; Scale bar, 5μm*.

Cdc15 promotes cytokinetic events after ring assembly, such as ring stabilization and septum ingression. Temperature sensitive *cdc15* mutants do not form a septum under restrictive conditions, and *cdc15* mutants lacking the SH3 domain show delays in the localization of the septum synthesizing enzyme Bgs1 (Arasada and Pollard, 2014; Cortes et al., 2015). In agreement with the fact that Cdc15 levels during maturation did not increase in CK666 treated cells (Fig. 5C), we find that under these conditions onset of ring constriction and septum ingression was also inhibited (Fig. 5E). CK666 treated cells were able to assemble an actomyosin ring as shown by Rlc1-tdTomato, but failed to recruit Bgs1-GFP (Fig. 5E). In these cells we observed an initial faint signal of Bgs1-GFP (Fig. 5E, green box) after ring assembly but the intensity of this signal did not increase with time and the rings did not constrict (Fig. 5E). Cells that had already recruited Bgs1 to the ring and initiated constriction prior to treatment were able to complete the process (Supplementary material Fig. S2B). CK666 treatment also inhibited cell separation at the end of cytokinesis (Supplementary material Fig. S2C). Thus after ring assembly, endocytic events dependent on the Arp2/3 complex contribute to ring constriction and septum ingression, likely through localization of Cdc15 and Bgs1 to the division site.

### Gef1 controls Cdc15 localization to the endocytic patches

We have shown that *cdc12Δ503* mutants exhibit lower levels of Cdc15 at the ring. Further, the *gef1Δcdc12Δ503* mutant displays uneven Cdc15 distribution along the ring. Therefore, we hypothesize that Gef1 regulates Cdc15 distribution along the ring. Cdc15 localizes to the ring during cytokinesis and to the endocytic patches during interphase. In cytokinesis, endocytosis initiates at the division site during the maturation phase. Previous reports indicate that in *cdc15* temperature sensitive mutants, endocytic patches do not appear at the division site under restrictive condition (Balasubramanian et al., 1998; Chang et al., 1996). Our data indicates that inhibiting endocytic patch formation by CK666 treatment prevents Cdc15 recruitment to the ring during maturation (Fig. 5C,D). Thus we investigated Cdc15 localization at the endocytic patches in gef *1Δcdc12Δ503* mutants. Since Cdc15 is a highly abundant protein in the cell, it is not possible to resolve Cdc15-GFP patches at the division site from proteins organized within the ring. To overcome this problem, we analyzed Cdc15-GFP patches at the cell tips during interphase. We find that cortical patches in gef *1Δ* and gef *1Δcdc12Δ503* mutants exhibit higher Cdc15-GFP intensity compared to *gef1*^+^*cdc12*^+^ cells (Fig. 6A, B). We show that compared to *gef1*^+^cdc12^+^*cells*, the mean intensity of Cdc15-GFP in cortical patches was higher by ~30% in *gef1Δ* (p<0.0001), ~14% in *cdc12Δ503* cells (p<0.05) and ~47% in *gef1Δcdc12Δ503* mutants, (p<0.0001; Fig. 6A,B). In addition, we found that in *gef1Δ* mutants, the number of Cdc15-GFP labelled patches increased at the cell tips. While *gef1*^+^*cdc12*^+^ and *cdc12Δ503* cells show an average of 11 patches per cell, *gef1Δ* and *gef1Δcdc12Δ503* mutants display 15 patches per cell (Fig. 6C).

**Figure 6.**
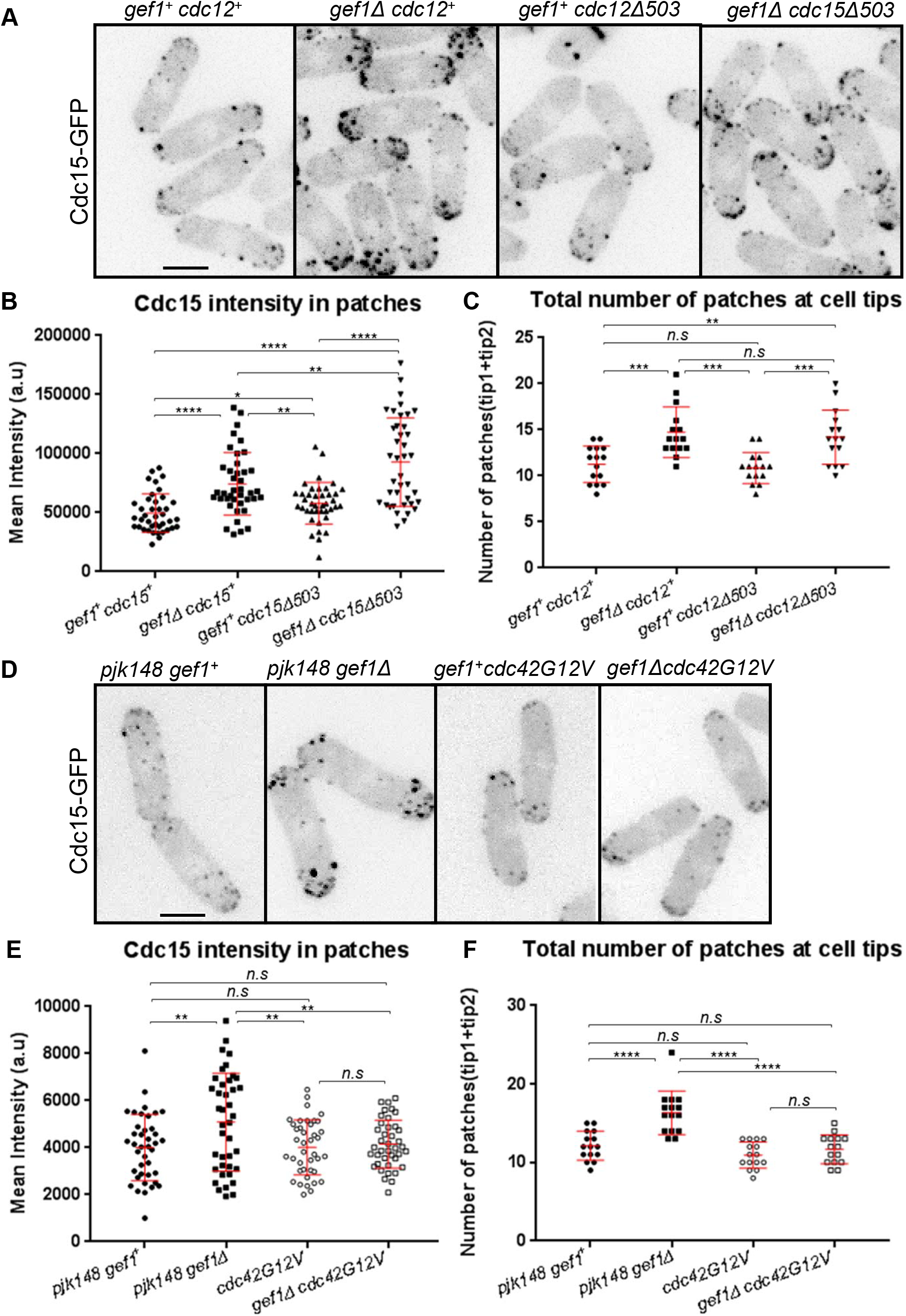
Gefl regulates Cdc15 localization to cortical patches. **(A)**. Cdc15-GFP labelled cortical patches in indicated strains. **(B).** Quantification of Cdc15-GFP mean fluorescence intensity in cortical patches in strains as indicated **(C).** Quantification of the number of Cdc15-GFP labelled cortical patches in interphase cells in strains as indicated. **(D).** Cdc15-GFP labelled cortical patches in *gef1*^+^ and *gef1Δ* cells transformed with the empty vector pJK148 or expressing constitutively active Cdc42, cdc42G12V. **(E).** Quantification of Cdc15-GFP mean fluorescence intensity in cortical patches in the strains represented in D. **(F).** Quantification of the number of Cdc15-GFP labelled cortical patches in interphase cells in strains represented in D. *n.s* −not significant; *\*p≤0.05*; \*\**p<0.01*; \*\*\**p<0.001*; \*\*\*\**p<0.0001; Scale Bar, 5μm; error bars, standard deviation*.

We asked whether Gef1 regulates Cdc15 localization to endocytic patches via activation of Cdc42. To test this, we expressed the constitutively active allele, *cdc42G12V* under a medium strength thiamine-repressible promoter *nmt41* in *gef1+* and *gef1Δ* cells. We have previously shown that under moderate expression levels, *cdc42G12V* rescues the delay in onset of ring constriction in *gef1Δ* cells (Wei et al., 2016). Here, *gef1+* and *gef1Δ* cells bearing the empty pjk148 vector serve as experimental controls. Again, we find that in *gef1Δ* cells bearing the control vector, Cdc15-GFP mean intensity was ~21 % higher than that of *gef1+* cells (p<0.01). In *gef1Δ* cells expressing moderate levels of *cdc42G12V*, Cdc15-GFP mean intensity is restored to those observed in *gef1+* cells bearing the control vector or expressing *cdc42G12V* (Fig. 6D, E). Furthermore, both *gef1+* and cdc42G12V expressing *gef1Δ* cells had an average of 12 patches per interphase cell as opposed to 16 patches in *gef1Δ* cells expressing the control vector (Fig. 6D, F). Thus, expression of *cdc42G12V* rescued the defects in Cdc15 patch intensity and numbers in *gef1Δ* mutants. These data indicate that Gef1 contributes to Cdc15 localization to the patches through Cdc42 activation.

### Model to define uniform protein distribution along an actomyosin ring-membrane interface

Our data show non-concentric furrowing in cells that display an irregular distribution of Cdc15 along the ring during maturation. To explore potential mechanisms governing how Cdc15 is recruited and distributed at the actomyosin ring during maturation, we developed a mathematical model. We find that Arp2/3-mediated actin nucleation is required for Cdc15 recruitment to the actomyosin ring after assembly. Furthermore, *gef1Δcdc12Δ503* cells display irregular Cdc15 distribution along the ring and non-concentric furrow formation. It has been shown that Cdc15 is recruited to endocytic patches immediately after initiation of actin nucleation. As the patch size increases, Cdc15 intensity at the patch also increases. However, Cdc15 does not internalize with the endocytic patch but is removed from the cortex by an unknown mechanism. Our model posits that endocytic patches are able to recruit Cdc15 to the vicinity of the assembled actomyosin ring. However, as the patches internalize, the Cdc15 associated with each patch remains at the cortex and is integrated into the ring. As endocytosis continues at the division site, Cdc15 recruitment persists and its levels increase at the actomyosin ring.

In the mathematical model, we define biophysical rules for the behavior of Cdc15 and characterize its spatial distribution as a function of time during ring maturation (Fig. 7A). The model is stochastic in nature and considers the behavior of individual Cdc15 proteins. The proteins can diffuse on the ring and associate with other Cdc15 molecules to form clusters, which we assume have a small diffusion coefficient and are effectively immobile. The population of Cdc15 increases over time due to recruitment from endocytic patches. Because Cdc15 proteins associated with an endocytic patch are spatially localized, they are recruited to the ring in a spatially correlated manner. Thus, if only a few large patches of Cdc15 are recruited, and the proteins do not spread around the ring, this can result in irregular protein distribution.

**Figure 7.**
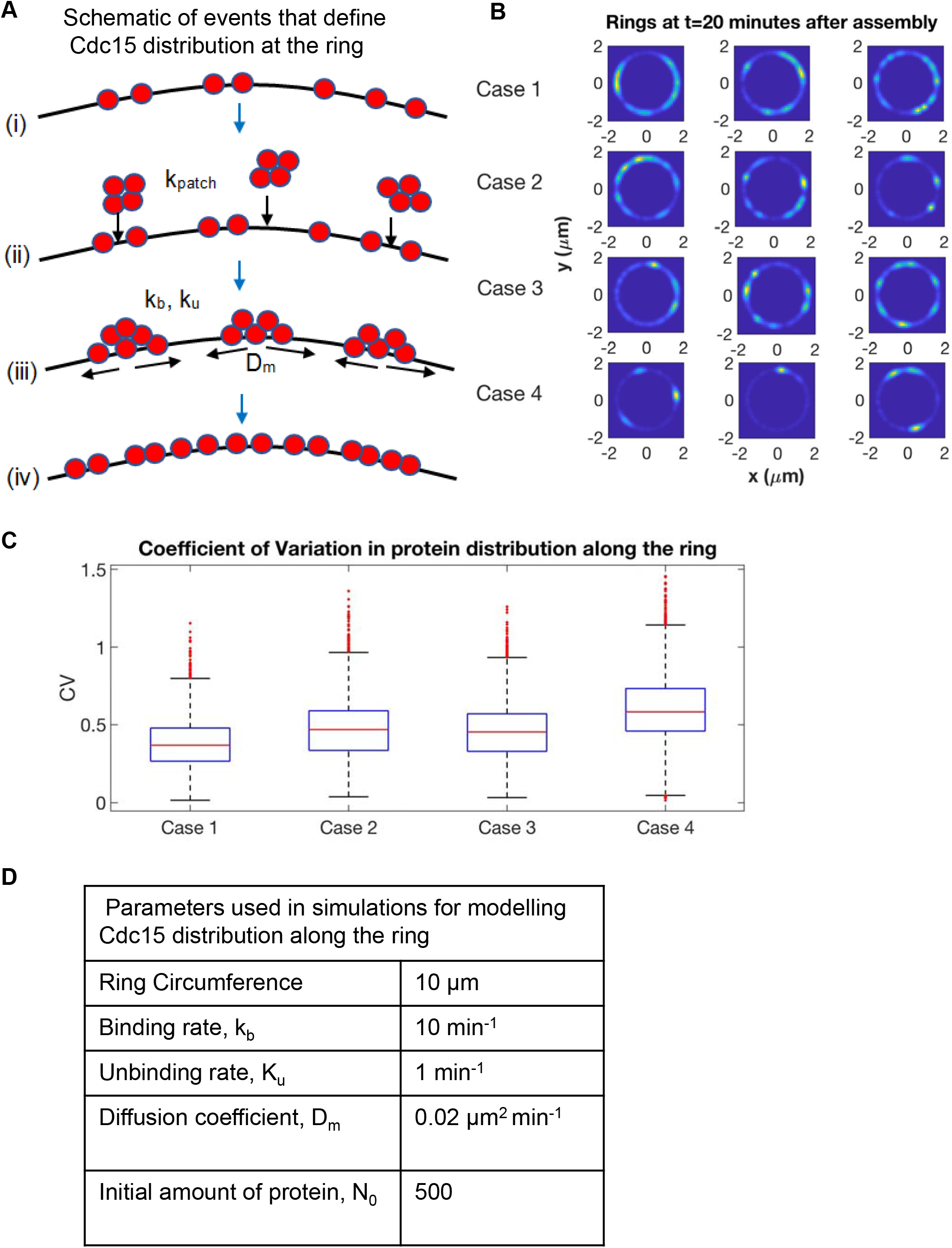
A mathematical model defines Cdc15 protein distribution along the ring during cytokinesis. **(A)**. Schematic of model (from top down): **(i).** A small initial population of Cdc15 proteins (red) are randomly distributed on the ring. **(ii).** Patches of Cdc15 proteins (from endocytic patches) associate to random locations on the ring at rate k_patch_. **(iii).** Proteins can bind to (rate k_b_) and unbind from (rate k_u_) each other and diffuse along the ring with diffusion coefficient D_m_. (iv). As time progresses, proteins accumulate and become distributed along the ring. **(B).** The spatial distribution of Cdc15 along the ring at t=20 min. Three independent simulated rings are shown for each of the 4 simulated cases. **(C).** Box plots of the coefficient of variation (CV) for 10,000 simulations of each of the four simulated cases. For each box, the central (red) mark is the median, the edges of the box are the 25^th^ and 75^th^ percentiles, and the whiskers extend to the most extreme data points not considered outliers. Outliers are plotted individually. The differences between all pairs of cases are significant, with p-values all below 4 × 10^-5^. **(D).** Parameter values used in all ring simulations. The number of proteins was obtained from previously published work (*Wu and Pollard, 2005*).

The model is parameterized by the rate at which endocytic patches are internalized near the ring (k_patch_), the number of Cdc15 molecules per patch (N_patch_), the total number of proteins in the ring at the end of maturation (N_total_), the diffusion coefficient of Cdc15 on the ring (D_m_), and the rates of binding to (k_b_) and unbinding from (k_u_) clusters. Details of the model can be found in the Supplementary Information. We investigated different sets of parameters representing our experimental observations in *gef1*+*cdc12*+, *gef1Δ, cdc12Δ503*, and *gef1Δcdc12Δ503* cells (Case1-4 respectively, Supplementary Figure S3). For each set, we performed computer simulations to generate multiple independent trajectories, each of which represents the spatiotemporal evolution of Cdc15 in the ring of a single cell. Figure 7B shows snapshots from simulations depicting the spatial distribution of Cdc15 on the ring for different genotypes (cases). We find that Cdc15 distribution in our simulated rings were uniform for cases 1-3, representing *gef1+cdc12+, gef1Δ*, and *cdc12Δ503* cells. However, Case 4, representing *gef1Δcdc12Δ503* cells, showed irregular Cdc15 distribution (Fig. 7B). We further verified this by measuring the coefficient of variation (CV) for 10,000 simulated rings for each case (Fig. 7C). We find that Case 1, simulating *gef1+cdc12+* conditions, has the lowest median CV and Case 4, simulating *gef1Δcdc12Δ503* conditions, has the largest median value with the distribution most prominently shifted toward larger values of CV. Because of the large number of simulations, differences between all distributions are statistically significant. Our simulations reveal that similar to experimental observations in *gef1Δcdc12Δ503* cells, Cdc15 distribution along the ring is irregular under conditions in which the number of molecules in each patch is increased and the rate of patch association is decreased. Thus, our model indicates that the distribution of Cdc15 along the ring is impacted by the patch association rate and the number of molecules in each patch.

## DISCUSSION

In fission yeast, coordinating ring constriction and septum ingression lead to concentric furrow formation. Here we report that the *gef1Δcdc12Δ503* mutant displays non-concentric furrow formation and irregular rate of constriction along the ring suggesting a disruption in the coordination of these processes. These mutants display an uneven distribution of the F-BAR protein Cdc15 at the ring such that sections of the ring constricting faster exhibit higher levels of the protein. Since previous reports show that Cdc15, a ring component, promotes septum ingression, defining what determines uniform Cdc15 distribution along the ring will provide insights into how ring constriction and septum ingression are coordinated. Our data indicate that trafficking events after ring assembly determine how Cdc15 is uniformly organized at the ring.

While Cdc15 distribution at the ring was irregular in cells displaying non-concentric furrow formation, we did not see any defects in F-actin organization or type II myosin distribution. This suggested that while the assembled rings themselves were spatially uniform, Cdc15 localization was not. This indicates that factors other than the assembled ring determine how proteins are organized along the ring. We find that Cdc15 dynamics are slower in *cdc12Δ503* and *gef1Δcdc12Δ503* mutants. Further, Cdc15 levels at the ring in late anaphase are lower in *cdc12Δ503* and *gef1Δcdc12Δ503* mutants. Our data suggests that the activated formin mutant influences Cdc15 localization to the ring. We posit that low levels of Cdc15 at the ring contributes to its irregular distribution and subsequent non-concentric furrow formation. Previous reports as well as our findings have shown that *cdc12Δ503* (Coffman et al., 2013) and *gef1Δcdc12Δ503* mutants display increased number of actin cables. Given that actin contributes to membrane trafficking events, it is possible that defects in membrane trafficking result in reduced Cdc15 levels at the ring in these mutants. This is supported by our observation that Cdc15 recruitment to the ring during maturation requires Arp2/3 complex dependent endocytic patch formation. This suggests that endocytosis is required for Cdc15 recruitment to the ring during maturation. While the rings in both *cdc12Δ503* and *gef1Δcdc12Δ503* mutants display decreased levels of Cdc15, only *gef1Δcdc12Δ503* mutant rings display irregular Cdc15 distribution and non-concentric furrow formation. This reveals a role for Gef1 in Cdc15 distribution along the ring. We show that Gef1 limits the levels of Cdc15 in individual endocytic patches at the cortex. Both *gef1Δ* and *gef1Δcdc12Δ503* mutants show increased levels of Cdc15 at the individual endocytic patches at the cortex.

Together our data indicates that Cdc15 distribution along the ring is irregular under conditions where Cdc15 displays slower dynamics at the ring and increased levels in individual cortical patches. We propose a model to explain how Cdc15 is organized at the ring after assembly and under what conditions this organization is disrupted. In our model, Cdc15 is incorporated into the ring from endocytic patches when endocytosis is initiated near the assembled actomyosin ring. Since Cdc15 remain in the patches until internalization, it is possible that its incorporation from the patches can be delayed if patch internalization itself is delayed. In addition, the amount of Cdc15 incorporated from each endocytic patch also influences its distribution. Under conditions where the number of molecules in each patch is high and the rate of patch incorporation is slow, Cdc15 distributes irregularly at the ring. This suggests that the rate of endocytosis at the division site and the amount of proteins in these patches together determine how Cdc15 is spatially distributed at the ring.

Why is uniform Cdc15 distribution needed for concentric furrow formation? In both *gef1* and *cdc15* mutants, Bgs1 localization is delayed at the ring with a corresponding delay in onset of furrow formation (Arasada and Pollard, 2014; Cortes et al., 2015; Wei et al., 2016). It is possible that Gef1 via Cdc42 activation regulates Cdc15 organization to ensure timely Bgs1 localization and onset of furrow formation. Cdc15 is an essential protein that localizes to the division site just before the onset of ring assembly (Wu et al., 2003). While Cdc15 supports actomyosin ring assembly, it plays a more important role in septum ingression and ring constriction (Arasada and Pollard, 2014; Cortes et al., 2015; Ren et al., 2015; Roberts-Galbraith et al., 2009). The SH3 domain of Cdc15 acts as a scaffold that binds several proteins and maintains them at the ring (Cortes et al., 2015; McDonald et al., 2017; Ren et al., 2015; Roberts-Galbraith et al., 2009). In addition, it also interacts with and localizes the GTPase Rho1 GEF, Rgf3 (Ren et al., 2015). Details of how Cdc15 promotes septum ingression remain unclear. One potential mechanism could be through the activation of Rho1, by localization of its GEF,Rgf3 which serves as the regulatory subunit for glucan synthases such as Bgs1 and Bgs4 (Arellano et al., 1996; Morrell-Falvey et al., 2005; Mutoh et al., 2005). In the absence of Cdc15 interaction, Rgf3 likely fails to localize and activate Rho1 at the ring and this in turn could fail to activate the glucan synthases. We postulate that sections of the ring with decreased Cdc15, fail to spatially activate the glucan synthases and thus display slower rates of ring constriction and septum ingression. Proper organization of Cdc15 along the ring is therefore important for uniform septum ingression in coordination with the actomyosin ring. Further investigations will determine how Gef1 regulates Cdc15. Gef1 localizes to the actomyosin ring and endocytic events occur at the vicinity of the ring. Thus, Gef1 and endocytic events at the actomyosin ring coordinate septum ingression and proper furrow formation.

Actomyosin ring-based cytokinesis is observed mainly in animal and fungal cells and the nature of furrow formation in these systems differ according to the cell type. Whereas in fungal cells cytokinesis involves coordination of actomyosin ring constriction and septum ingression, in animal cells ring constriction is coordinated with membrane expansion (Prekeris and Gould, 2008). In certain animal cells such as embryos and polarized epithelial cells, furrow formation does not occur in a concentric manner (Maddox et al., 2007). These cells display asymmetric furrow formation in which the ring unilaterally constricts to one side in a non-concentric manner (Maddox et al., 2007). This manner of constriction likely maintains cell-cell contacts in epithelial cells and supports rapid cell division during embryonic development (Dorn et al., 2016; Maddox et al., 2007). It has been reported that septin and anillin scaffold proteins are required for non-concentric furrowing (Maddox et al., 2007). A recent model indicates that a non-concentric furrow can form even in the absence preexisting cues due to feedback between membrane curvature and cytoskeletal rearrangements (Dorn et al., 2016). Proper furrow formation in fission yeast has been described in a mathematical model which indicates that septum deposition is a mechanosensitive process and that actomyosin ring tension and curvature drives this process (Thiyagarajan et al., 2015). An impaired actomyosin ring results in irregular septum ingression and furrow formation. Our data provide a mechanistic insight into how the actomyosin ring acts as a landmark where proteins are organized to promote the septum ingression and furrow formation.

## MATERIALS AND METHODS

### Strains and Cell culture

Strains used in this study are listed in Supplemental Table S1. All S. *pombe* strains used in this study are isogenic to PN972. Unless otherwise mentioned cells were cultured in yeast extract (YE) medium and grown exponentially at 25°C. Genetic manipulation of strains were carried out using standard techniques (Moreno et al., 1991). Cells were grown exponentially for at least 3 rounds of eight generations each assay.

### Microscopy

Image acquisition was performed at room temperature (23-25°C) with the VT-Hawk 2D-array scanning confocal microscope (Visitech intl., Sunderland, UK) using an Olympus IX-83 inverted microscope with a 100x/numerical aperture 1.49 UAPO lens (Olympus, Tokyo, Japan). For still images, cells were mounted directly onto glass slides with a #1.5 coverslip (Fischer Scientific, Waltham, MA) and imaged promptly. For all z-series, images were acquired with depth interval of 0.4um. In time-lapse image acquisition, cells were placed in a 3.5-mm glass-bottom culture dish and covered with YE medium with 0.6% agar. Ascorbic acid (100uM vitamin C) was added to cell culture to minimize fluorescence toxicity, as previously reported (Wei et al., 2017).

Images were acquired with MetaMorph (Molecular Devices, Sunnyvale, CA). For the fluorescence recovery after photobleaching experiments the SP8 confocal microscope with 63x 1.4NA oil objective (Leica Microsystems) was used. Images were acquired with the Leica Application Suite X software. All images were analyzed with Image J (National Institutes of Health, Bethesda, MD). Statistical significances between two groups of cells were determined by p-values from the Student’s t-test.

### Phalloidin staining

The actin cytoskeleton was stained as described previously (Das et al., 2009; Pelham and Chang, 2002). We fixed exponentially growing cells with 3.5% formaldehyde (Sigma Aldrich) and incubated at room temperature (23-25°C) for 10 minutes. The fixed cells were washed with PM buffer (35 mM KPO4, pH 6.8, 0.5 mM MgSO4) and permeabilized with 1% Triton in PM buffer. Finally, cells were washed again in PM buffer and then stained with Alexa-Fluor Phalloidin at room temperature for 30-45 minutes.

### Cdc15 patch analysis

For Cdc15-GFP cortical patch analysis we acquired images in z-series. The images were SUM projected and the circumference of each patch was traced using the freehand tool in ImageJ. The mean fluorescence intensity in each patch was measured. A region outside the cell was used for background subtraction. Mean fluorescence intensity was reported in arbitrary units (a.u). To count the number of patches at the cell tips, we analyzed z-series of images. We counted the number of patches for each cell. To eliminate any error due to overlapping patches we scrolled through each Z-step in ImageJ, using the labelling tool to identify each patch.

### Fluorescence recovery after photobleaching (FRAP)

Cells with an actomyosin ring in the maturation phase were first focused at the medial Z plane. At the medial plane the Cdc15-GFP labelled ring appears as two spots on either edge of the cell cortex. One such spot was marked as the region of interest and 4 images were acquired prebleach. The region of interest was then bleached for 10 seconds and images were then acquired every 3 seconds for 50-100 images. The non-bleached spot for the same ring was also analyzed to account for loss of signal due to photobleaching over time. A cell free region of the image was used for background subtraction. Intensity values for each region of interest (ROI) was corrected for loss of signal over time and background. The intensity values were then normalized against the mean pre-bleach value to determine the recovery fraction over time. For each genotype at least 10 rings were analyzed. The average corrected and normalized intensity values were then plotted as fraction of recovery over time. The data was then fitted to a single-exponential curve of the functional form *f*(*x*) = *m*_1_ + *m*_2_*e*^−*m*_3_*x*^, where *m*_3_ is the off rate and *m*_1_ is the plateau of recovery. The half-life, *t*_1/2_, was then computed as 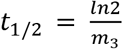. Biphasic recovery curves were then constructed using a double-exponential form of the curve-fitting function, *f*(*x*) = *m*_1_ + *m*_2_*e*^−*m*_3_*x*^ + *m*_4_*e*^−*m*_5_*x*^.

### CK666 treatment

Cells were treated with 100uM CK666 (Sigma Aldrich, SML006-5MG) in dimethyl sulfoxide (DMSO by Sigma-Aldrich, D8418-250ML). Control cells were treated with 0.1%DMSO in YE medium. For time-lapse images the agar pads (Wei et al., 2017) were also treated with 100uM CK666 or DMSO.

### Expressing constitutively active Cdc42

The *cdc42G12V* fragment was cloned into the pJK148 vector under the thiamine-repressible promoter nmt41 and integrated into the genome of *gef1+ and gef1Δ* cells as previously described in Wei *et al*., 2016. To ensure low expression levels of cdc42G12V cells were grown in YE medium at 25°C. The experimental controls were *gef1+ and gef1Δ* cells transformed with the empty pJK148 vector.

## ACKNOWLEDGEMENTS

We thank B. McKee, B. Hercyk and J. Rich for critically reviewing our manuscript; and J.Q. Wu, K. Gould, and P. Perez for providing strains.

